# Cardiomyocyte Targeting Peptide to Deliver Amiodarone

**DOI:** 10.1101/2023.05.10.540206

**Authors:** Maliha Zahid, Beth Weber, Ray Yurko, Kazi Islam, Vaishavi Agrawal, Jack Lopuszynski, Hisato Yagi, Guy Salama

**Affiliations:** Dept. of Cardiovascular Diseases, Mayo Clinic, Rochester, MN; Pittsburgh Heart, Lung, Blood, and Vascular Medicine Institute and Division of Cardiology, Department of Medicine, University of Pittsburgh School of Medicine and University of Pittsburgh Medical Center, Pittsburgh, PA, USA; Peptide Synthesis Facility, University of Pittsburgh, Pittsburgh, PA; Dietrich School of Arts and Sciences, University of Pittsburgh, Pittsburgh, PA; Burnett School of Biomedical Sciences, University of Central Florida, Orlando, FL; Dept. of Developmental Biology, University of Pittsburgh, Pittsburgh, PA

## Abstract

**Background:** Amiodarone is underutilized due to significant off-target toxicities. We hypothesized that targeted delivery to the heart would lead to lowering of dose by utilizing a cardiomyocyte targeting peptide (CTP), a cell penetrating peptide identified by our prior phage display work.

**Methods:** CTP was synthesized thiolated at the N-terminus, conjugated to amiodarone via Schiff base chemistry, HPLC purified and confirmed with MALDI/TOF. Stability of the conjugate was assessed using serial HPLCs. Guinea pigs (GP) were injected intraperitoneally daily with vehicle (7 days), amiodarone (7 days; 80mg/Kg), CTP-amiodarone (5 days;26.3mg/Kg), or CTP (5 days; 17.8mg/Kg), after which GPs were euthanized, hearts excised, perfused on a Langendorff apparatus with Tyrode’s solution and blebbistatin (5μM) to minimize contractions. Voltage (RH237) and Ca^2+^-indicator dye (Rhod-2/AM) were injected, fluorescence from the epicardium split and focused on two cameras capturing at 570–595nm for cytosolic Ca^2+^ and 610–750nm wavelengths for voltage. Subsequently, hearts were paced at 250ms with programmed stimulation to measure changes in conduction velocities (CV), action potential duration (APD) and Ca^2+^ transient durations at 90% recovery (CaTD_90_). mRNA was extracted from all hearts and RNA sequencing performed with results compared to control hearts.

**Results:** CTP-amiodarone remained stable for up to 21 days at 37°C. At ∼1/15^th^ of the dose of amiodarone, CTP-amiodarone decreased CV in hearts significantly compared to control GPs (0.92±0.05 vs. 1.00±0.03m/s, p=0.0007), equivalent to amiodarone alone (0.87±0.08ms, p=0.0003). Amiodarone increased APD (192±5ms vs. 175±8ms for vehicle, p=0.0025), while CTP-amiodarone decreased it significantly (157±16ms, p=0.0136) similar to CTP alone (155±13ms, p=0.0039). Both amiodarone and CTP-amiodarone significantly decreased calcium transients compared to controls. CTP-amiodarone and CTP decreased CaTD_90_ to an extent greater than amiodarone alone (p<0.001). RNA-seq showed that CTP alone increased the expression of DHPR and SERCA2a, while decreasing expression of proinflammatory genes NF-kappa B, TNF-α, IL-1β, and IL-6.

**Conclusions:** Our data suggests that CTP can deliver amiodarone to cardiomyocytes at ∼1/15^th^ the total molar dose of amiodarone needed to produce comparable slowing of CVs. The ability of CTP to decrease AP durations and CaTD_90_ may be related to its increase in expression of Ca-handling genes, and merits further study.

## Introduction

Atrial fibrillation (afib) is the most common arrhythmia in adults with an estimated prevalence of 46.3 million worldwide^1^. It affects 1-2% of adults in the United States, with an estimated 5.7 million patients with afib^2^. Data from the Framingham heart study has documented an over three-fold increase in afib prevalence over the last 50 years^3^. These already high rates are estimated to double by year 2050 due to an aging population, and increased prevalence of risk factors like coronary artery disease, hypertension, obesity, and congestive heart failure^3^. Superiority of rhythm over rate control in afib has been demonstrated in multiple randomized control trials^4-6^. Although amiodarone is the most efficacious anti-arrhythmic drug with demonstrated superiority over multiple other anti-arrhythmics like propafenone^7,8^, sotalol^7,9,10^, and dronedarone^11,12^, it is grossly underutilized and remains a second-line therapy for maintenance of normal sinus rhythm in patients with afib. This is due to amiodarone being a highly lipophilic drug with a large volume of distribution^13^, a long half-life, uptake by non-cardiac tissues like lungs, liver, thyroid, skin and ocular tissue, leading to multiple off-target, potentially life-threatening toxicities thus limiting its use to patients over 75 years of age, or patients with limited life expectancies^14^. Additionally, amiodarone is the drug of choice for ventricular tachycardia^15,16^, as well as for reducing defibrillator shocks in patients with cardiomyopathies who have an implanted automatic intravascular defibrillation^16,17^. Hence, amiodarone is a classic demonstration of the Achilles heel of both cardiac diagnostics and therapeutics suffering from a lack of cardiac specific vector(s) and lack of targeted delivery.

The cell plasma membrane is a semi-permeable barrier that is essential for cell integrity and survival but at the same time presents a barrier to delivery of cargo. Hence, the ability of certain proteins to cross cell membrane barriers was met with great enthusiasm. In 1988, two separate groups demonstrated the ability of trans-activator of transcription (Tat) protein of the human immunodeficiency virus (HIV) to enter cultured cells and promote viral gene expression^18,19^. Mapping of the domain responsible for this transduction ability led to the identification of the first cell-penetrating peptide, Tat, corresponding to the 11 amino acid basic domain of HIV-1 Tat protein. Subsequently, it was demonstrated that Tat fused to β-galactosidase and injected intra-peritoneally into mice was internalized into multiple cell types including liver, heart, lung, kidney, and even crossed the blood brain barrier delivering β-galactosidase in a functional form, highlighting the potential of these peptides as vectors^20^.

The ability of cationic or hydrophobic cell penetrating peptides to transduce a wide variety of tissue types *in vivo* limits their clinical utility because of their lack of cell specificity, leading to a higher chance of off-target, adverse side effects. Phage display using libraries of various lengths and various bacteriophage strains have been successfully utilized to identify tissue-specific penetrating peptides^21-25^. In our prior work, we employed a combinatorial *in vitro* and *in vivo* M13 phage display methodology^21^ to identify a 12-amino acid, synthetic, mildly basic, non-naturally occurring peptide (NH_2_-APWHLSSQYSRT-COOH) that we termed *C*ardiac *T*argeting *P*eptide (CTP), due to its ability to specifically target cardiomyocytes *in vivo* after a tail-vein injection in mice^21,26^. Our detailed bio-distribution studies show that peak uptake occurs as early as 15 minutes with complete disappearance of fluorescently labeled CTP by 6 hrs^27^. CTP does not cross the blood-brain barrier, is not taken up by striated muscle, spleen, lungs etc. CTP appears to be excreted predominantly by a renal, followed by hepatobiliary mechanism^27^. Our findings were replicated by at least three independent laboratories with published work showing that indeed CTP is cardiomyocyte specific^28-30^. It has been used to deliver photosensitizers to the cardiomyocytes for targeted ablation of afib in a sheep model, sparing damage to “innocent bystander” cardiac myofibroblasts and endothelial cells^28^. Additionally, exosomes labeled with CTP and loaded with anti-RAGE (*R*eceptor for *A*dvanced *G*lycation *E*nd-products) siRNA was able to ameliorate myocarditis in a rat model^30^. CTP is not limited to mice, as its efficacy as a vector has been demonstrated in sheep^28^ and rat models^30^ to deliver myriad cargoes. Indeed, our own lab showed that human heart tissue explanted from patients undergoing heart transplants could be successfully transduced with fluorescently labeled CTP with uptake of the peptide by normal cardiomyocytes while sparing the fibroblasts that make up the intervening scar tissue in these diseased hearts. Moreover, the uptake in cardiomyocytes was not due to an increase in plasma membrane permeability as demonstrated by a lack of uptake of Evans blue ^31^.

In the current manuscript, we present studies showing successful conjugation of amiodarone to the N-terminus of CTP via a disulfide linker. We tested our conjugate in guinea pigs (GPs) to compare changes in calcium handling equivalent to that seen with amiodarone but at markedly decreased (1/15^th^) total dose of amiodarone. A surprise finding was the additional effects on calcium transients that were attributed to CTP portion of the CTP-amiodarone conjugate and not amiodarone. This led us to perform gene expression studies which showed that CTP had additional salutary anti-inflammatory effects and changes in calcium handling genes. Our findings lead us to conclude that CTP is not simply an inert vector but has additional, potentially beneficial, biological effects on cardiomyocytes.

## Methods

### CTP Synthesis and Conjugation to Amiodarone

Solid phase peptide synthesis (SPPS) of CTP was accomplished on a Liberty CEM microwave synthesizer using Fmoc/tBu chemistry and Oxyma pure coupling protocols on Rink amide resin. Upon completion of CTP-peptide chain assembly, the free N-terminal amino group was manually conjugated to 3-(Tritylthio)propionic acid with TBTU/HOBt/DIPEA in DMF. The Thiol modified CTP peptide resin was then cleaved with Trifluoroacetic acid (TFA) + scavengers followed by isolation of the crude product by precipitation in Diethyl Ether (EtO2). Crude CTP-thiol was then purified by preparative C-18 RP-HPLC on a Waters Delta Prep 4000 chromatography system followed by Lyophilization. Purified CTP-thiol was then reacted with 2,2’-Dithiobis(5-nitropyridine) (DTNP) in 80% TFA(aq) for 15 minutes and then dried to a film using a stream of nitrogen. Dissolution of the 5-Nitro-2-Pyridinesulfenyl (pNpys) activated CTP analogue in 50%Trifluoroethanol/0.1% TFA was followed by preparative C-18 RP-HPLC purification on a Waters Delta Prep 4000 chromatography system and Lyophilization to a dry powder.

Amiodarone Hydrochloride was chloroalkylated at the tertiary nitrogen using 3-Chloro-1-propanethiol forming a thiolated quanternary ammonium salt. Amiodarone-thiol was then purified by preparative C-18 RP-HPLC purification on a Waters Delta Prep 4000 chromatography system and then Lyophilized. Purified amiodarone-thiol was then reacted with pNpys-CTP in ammonium acetate buffered solution at pH 4 for 2 hours. Preparative C-18 RP-HPLC purification on a Waters Delta Prep 4000 chromatography system was followed by Lyophilization to a dry powder. Analytical C-18 RP-HPLC characterization of Amiodarone-SS-CTP on a Waters Alliance chromatography system was performed followed by mass analysis on a Bruker UltraFlextreme MALDI TOF/TOF mass spectrometer to confirm the expected mass and purity of the final product.

### CTP Targeting of Cardiomyocytes

Wild-type Sprague Dawley rat heart was mounted in a Langendorff perfusion system and perfused with 50μl of a 1mM solution of Cy5.5 labeled CTP. After the bolus injection, heart was fixed in paraformaldehyde, embedded, cryosectioned with 7μm thick sections obtained, that were cross-stained with WGA-488 (wheat germ agglutinin labeled with Alexa fluoro dye 488) to stain for myofibroblasts and DAPI, a nuclear stain and confocal imaging performed.

### Conjugate Stability Studies

Freshly synthesized/purified CTP-amiodarone conjugate was HPLC characterized after lyophilization as baseline measure. A small amount (1mg/ml) was dissolved in the buffer used for injecting animals and placed in a water-bath at 37°C for upto 22 days. A small aliquot was taken at regular intervals and an HPLC run on Waters Delta Prep 4000 chromatography system.

### Guinea Pig Studies

All animal protocols were approved by the University of Pittsburgh’s institutional animal care and use committee prior to undertaking any experimentation. Adult, male, 6-week-old GPs (250-350gm) were injected intraperitoneally daily with vehicle (7 days), amiodarone (7 days; 80mg/Kg), CTP-Amiodarone (5 days; 26.3mg/Kg), or CTP (5 days; 17.8mg/Kg). At the end of the study, GPs were euthanized, hearts excised and perfused on a Langendorff apparatus with Tyrode’s solution containing (in mM): NaCl (130), KCl (4), MgSO_4_ (1.2), NaHCO_3_ (25), Glucose (5), CaCl_2_ (1.25), Mannitol (45) gassed with 95% O_2_ and 5% CO_2_, pH 7.0 at 37°C. Hearts were placed in a custom-designed chamber to subdue motion artifacts, along with a 15 minute blebbistatin (5 μM) in perfusate to minimize contractions. Bolus injections of voltage (RH237, 25μl of 2mg/mL dimethyl sulfoxide (DMSO)) and Ca^2+^-indicator dye (Rhod-2/AM, 200 μl of 2 mg/mL DMSO) were injected in the air-trap above the aortic cannula. Fluorescence from the epicardium was collected with a camera lens, split with a 570nm dichroic mirror and focused on two CMOS cameras (Sci-Media UltimaOne) capturing the fluorescence emission at 570– 595nm for cytosolic Ca^2+^ and 610–750nm wavelengths for voltage, as described in detail previously^32^. After resting sinus rhythm image acquisition, hearts were paced at 250ms with programmed stimulation to measure and compare conduction velocity and transient duration (90%) of various treatment groups.

### RNA Sequencing Studies

Libraries were generated with the Illumina Stranded mRNA Library Prep kit (Illumina: 20040534), according to the manufacturer’s instructions. RNA was assessed for quality using a Total RNA 15nt kit (Agilent: DNF-471-33) on an Advanced Analytical 5300 Fragment Analyzer. RNA concentration was quantified with a Qubit BR RNA assay kit (Invitrogen: Q10211) on a Qubit 4 (Invitrogen: Q33238). Briefly, 50 ng of input RNA was used for each sample. Following adapter ligation, 15 cycles of indexing PCR were completed, using IDT for Illumina RNA UD Indexes (Illumina: 20040553). Library assessment and quantification was done using Qubit 1x HS DNA (Invitrogen: Q33231) on a Qubit 4 fluorometer and a HS NGS Fragment kit (Agilent: DNF-474-1000). Libraries were normalized and pooled to 2nM by calculating the concentration based off the fragment size (base pairs) and the concentration (ng/μl) of the libraries. Sequencing was performed on an Illumina NextSeq 2000, using a P3 200 flow cell (Illumina: 20046812). The pooled library was loaded at 750 pM and sequencing was carried out with read lengths of 2x101 bp, with an average of ∼40 million reads per sample. Sequencing data was demultiplexed by Illumina the on-board DRAGEN FASTQ Generation software v3.8.4.

The raw RNA sequencing (RNA-Seq) paired-end reads for the guinea pig (GP) samples were processed through the Mayo RNA-Seq bioinformatics pipeline, RapMap version 1.0.0. Briefly, raw reads were trimmed with a base quality cut off of Phred score Q30 using Trimmomatic^33^. RapMap employs the very fast and accurate pseudo aligner and transcript assembler, Kallisto (https://pachterlab.github.io/kallisto/manual), to align the filtered reads to the reference GP genome Cavia_porcellus.Cavpor3.0. Kallisto was also used to calculate relative expression values across transcripts. Tximport^34^ was used to summarize transcript level counts at gene level. Finally, comprehensive quality assessment of the RNA-Seq samples was performed using MultiQC^35^. Results from all modules described above will be linked through a single html document and reported by the RapMap pipeline.

Using the gene-level counts from RapMap, genes differentially expressed between the Control and treated groups were assessed using the bioinformatics package edgeR 2.6.2 ^36^. Differentially expressed genes (DEGs) were reported along with their magnitude of change (log2 scale) and their level of significance (False Discovery Rate, FDR < 5%). DEGs were plotted using volcano plots and heatmaps using R Bioconductor packages ggplot2 (https://ggplot2-book.org/) and heat map (https://cran.r-project.org/web/packages/pheatmap/index.html) respectively. Canonical pathway analysis was performed using the Enrichr^37^ database. Pathways identified using KEGG were reported. R package pathview (https://pathview.r-forge.r-project.org/) was used to visualize the gene expression changes in pathways of interest.

### Statistical Analyses

All continuous variables generated on GP hearts were checked for normality with the skewness and normality test. Variables in each treatment group were compared to controls using an unpaired Student’s t-test. A two-tailed p-value of <0.05 was considered significant. Stata 16.1 (Stata Corporation, College Station, Texas) was used for these analyses.

## Results

### CTP Synthesis, Targeting, Conjugation to Amiodarone, and Conjugate Stability

This conjugation was accomplished with a disulfide bond between the quarternary amine salt of amiodarone and the N-terminus of CTP (Fig. 1). The rationale behind this approach was to create a conjugate that was stable in serum but once internalized into cardiomyocytes, the reducing intracellular enviornment would reduce the bond releasing amiodarone from CTP. This bond was stable for upto 22 days (longest time interval tested) at 37°C (Fig. 2). Rat hearts perfused with Cy5.5 labeled CTP showed robust uptake of CTP (red) by cardiomyocytes, and negligible co-localization with WGA-488, or uptake of CTP by myofibroblasts (Fig. 3).

**Figure 1:**
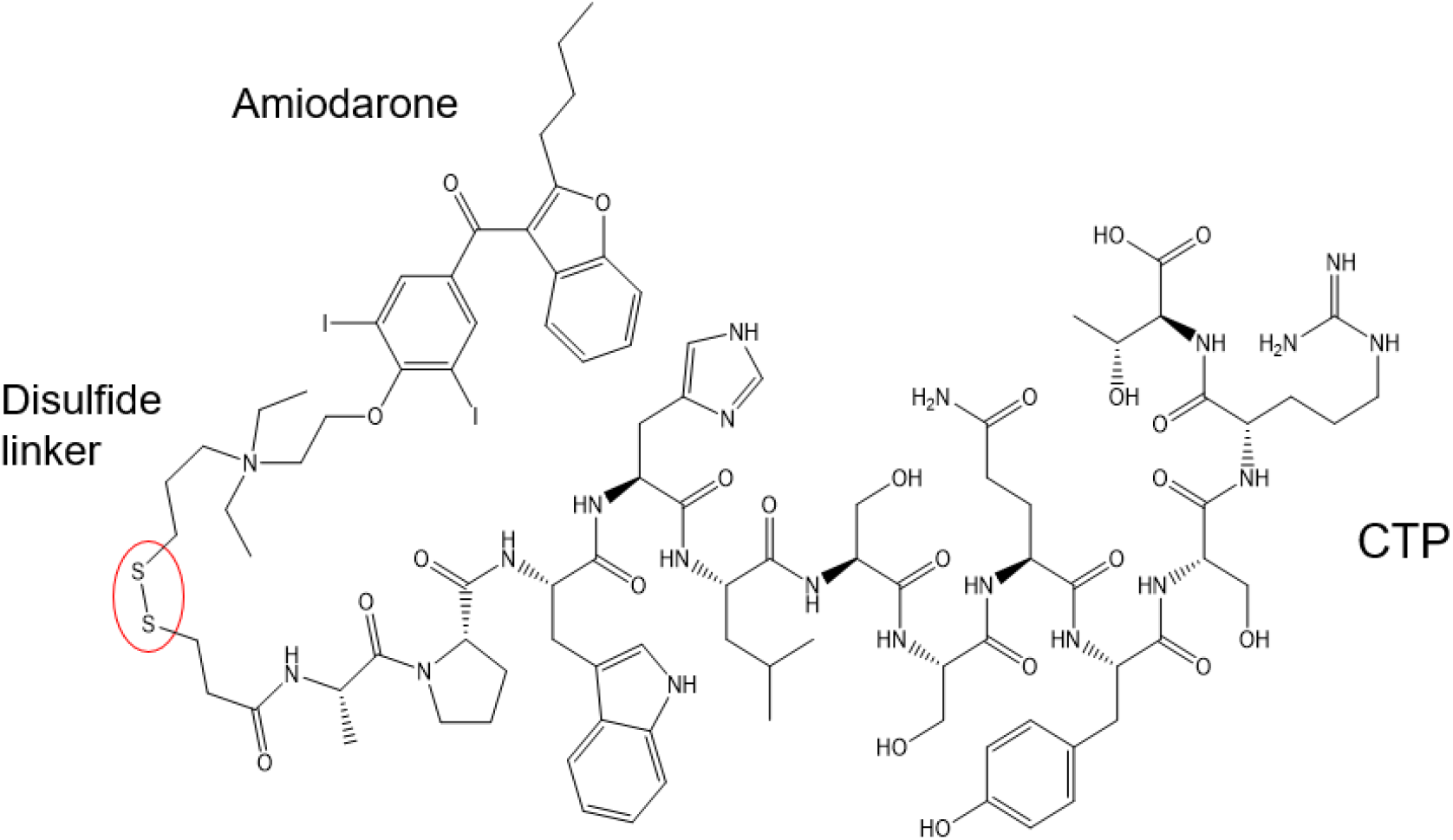
Schematic presentation of the CTP-amiodarone chemical structure. CTP is conjugated to amiodarone via a disulfide linker (circled in red).

**Figure 2:**
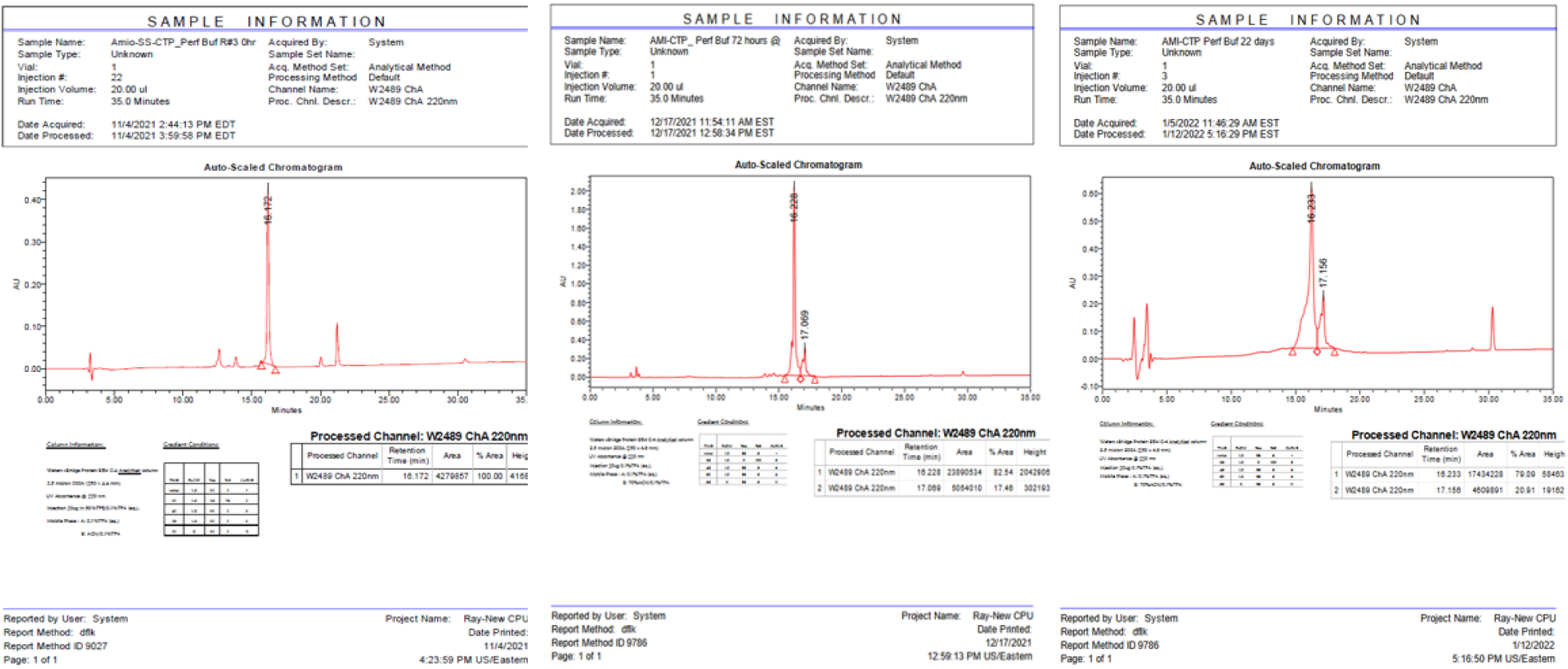
HPLC Tracing of the CTP-amio conjugate immediately upon synthesis and purification (a), after 72 hours of incubation (b), and after 22 days of incubation (c) at 37°C.

**Figure 3:**
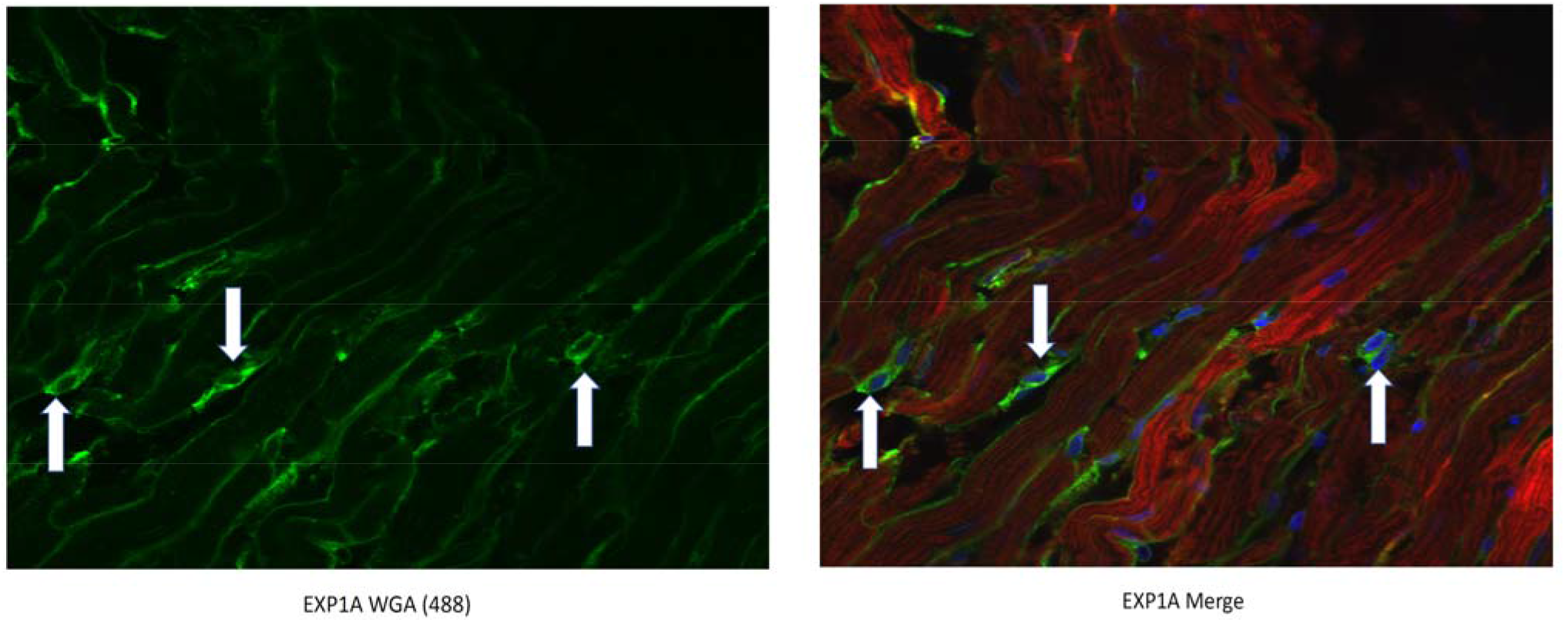
CTP-Cy5.5 transduces rat cardiomyocytes in a cardiomyocyte-specific manner. Myofibroblasts are stained green with WGA-488 and show no co-localization with the CTP-Cy5.5 (red) that is taken up exclusively by cardiomyocytes. Nuclei stained blue (DAPI).

### Guinea Pig Studies

Adult, male, GPs (250-350gm) were injected intraperitoneally with amiodarone (80mg/Kg; n=4) daily for 7 days, or CTP-amio (26.3mg/Kg; n=4) for 5 days or CTP alone (17.8mg/Kg; n=4) for 5 days, and compared to vehicle only injected GPs (n=10). At the end of the injection period, GPs were euthanized, hearts excised and perfused in a Langendorf apparatus. Hearts were labeled with a voltage-sensitive dye (RH 237) and a Ca^2+^ indicator (Rhod-2/AM) with a bolus injection in the perfusate, in air trap above the aortic cannula. Fluorescence images of the heart were focused on 2 CMOS camera, one to detect voltage, the other intracellular free Ca^2+^. Heart rate, action potentials (APs) and intracellular free-Ca^2+^ transients (CaT) weremeasured from 10,000 pixels of the voltage and Ca^2+^ CMOS cameras. Durations of APs and CaTs were calculated from the first derivative of the rise of the signals to their recovery to 90% of the baseline. Amiodarone injections significantly decreased heart rate during sinus rhythm, decreased the conduction velociy (CV) of the APs and CaT, increased AP and CaT durations compared to control hearts (Fig. 4a-e). The conjugate CTP-amio decreased CVs to values similar to amiodarone, and statistically significant decreases compared to control hearts. However, in contrast to amiodarone, CTP-amio increased heart rate, decreased APand CaT durations, effects that seem to be driven by the CTP component of the conjugate, because the conjugate alone, CTP decreased the AP and CaT duration (Fig. 4).

**Figure 4:**
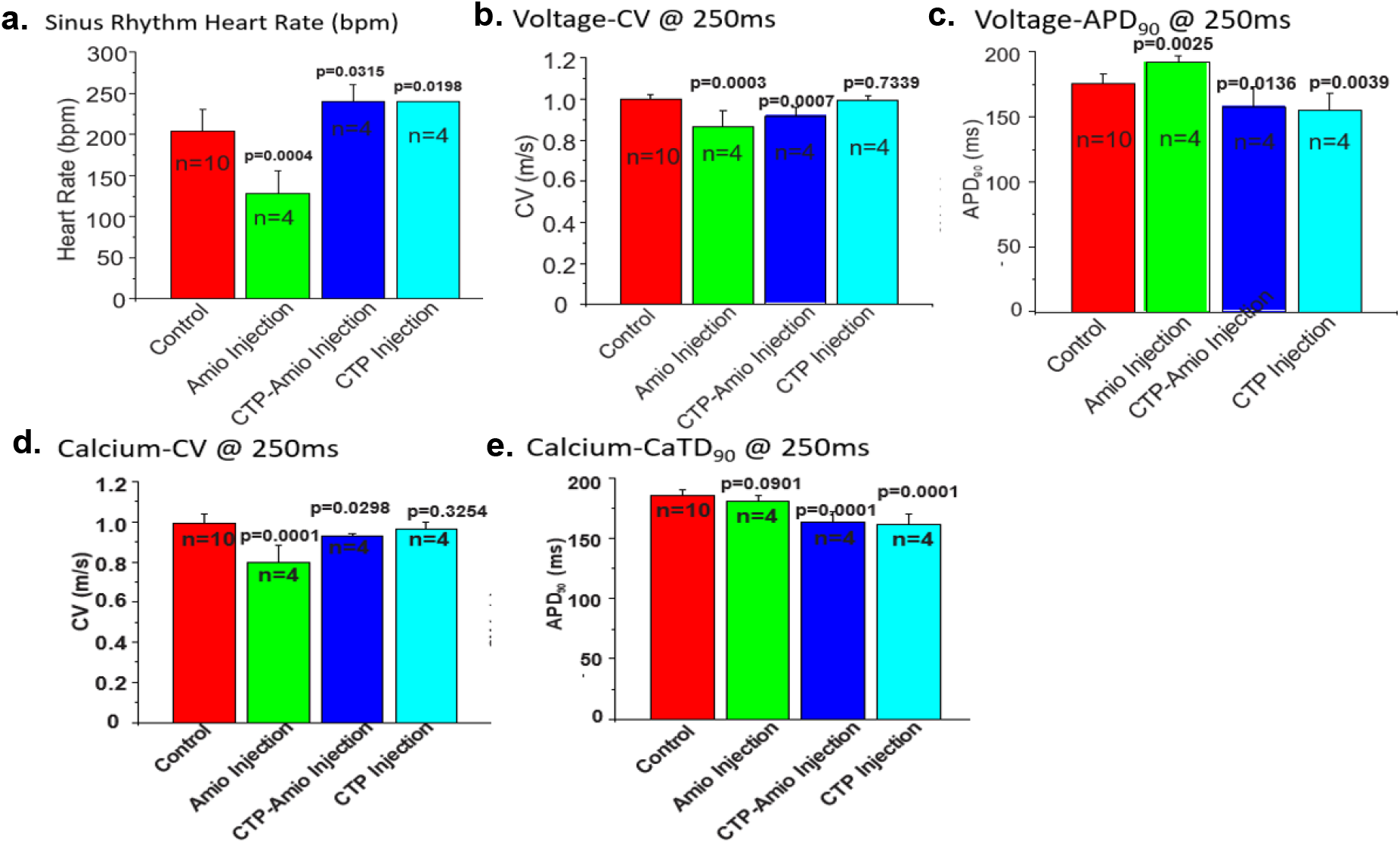
Comparison of heart rate (a), voltage conduction velocities (b), action potential duration (c), calcium conduction velocities (d) and CaT (e) between various treatment groups. All p-values are two-tailed, unpaired t-tests between each treatment group and controls.

### RNA Sequencing Studies

The mRNA was extracted from the 16 GP (4 controls, 4 amio, 4 CTP-amio, and 4 CTP only treated) hearts, sequenced with 20 million reads per sample, reads aligned to GP cDNA using Kalisto program for assessing degree of alignment (Supplemental Figure 1) which showed that ∼80% of the reads were successfully aligned (Supplemantal Figure 2). Principal componenet analysis showed good clustering of samples within groups and separation between groups (Fig. 5). Because of the unexpected effects of CTP alone, which was thought to be inert to cardiomyocytes, we now present the comparison of CTP treated hearts to control hearts, as well as amiodarone and CTP-amio treated hearts in the supplementary data (Supplemental Figures 3-5). In CTP-treated hearts compared to controls, there was an upregulation of α-adrenergic receptors and down-regulation of β-adrenergic receptors, possibly explaining the increase in resting heart rate in CTP (Fig. 6) and CTP-amio treated hearts (Supplemental Figure 6). Additionally, calcium handling channel genes, *DHPR* and *SERCA2a*, were significantly upregulated (Fig. 6), possibly explaining the decrease in late CaTs. Several pro-inflammatory genes, like *NF-κB, TNF-α, IL-1β*, and *COX2* were downregulated in CTP treated hearts as compared to control hearts (Fig. 7).

**Figure 5:**
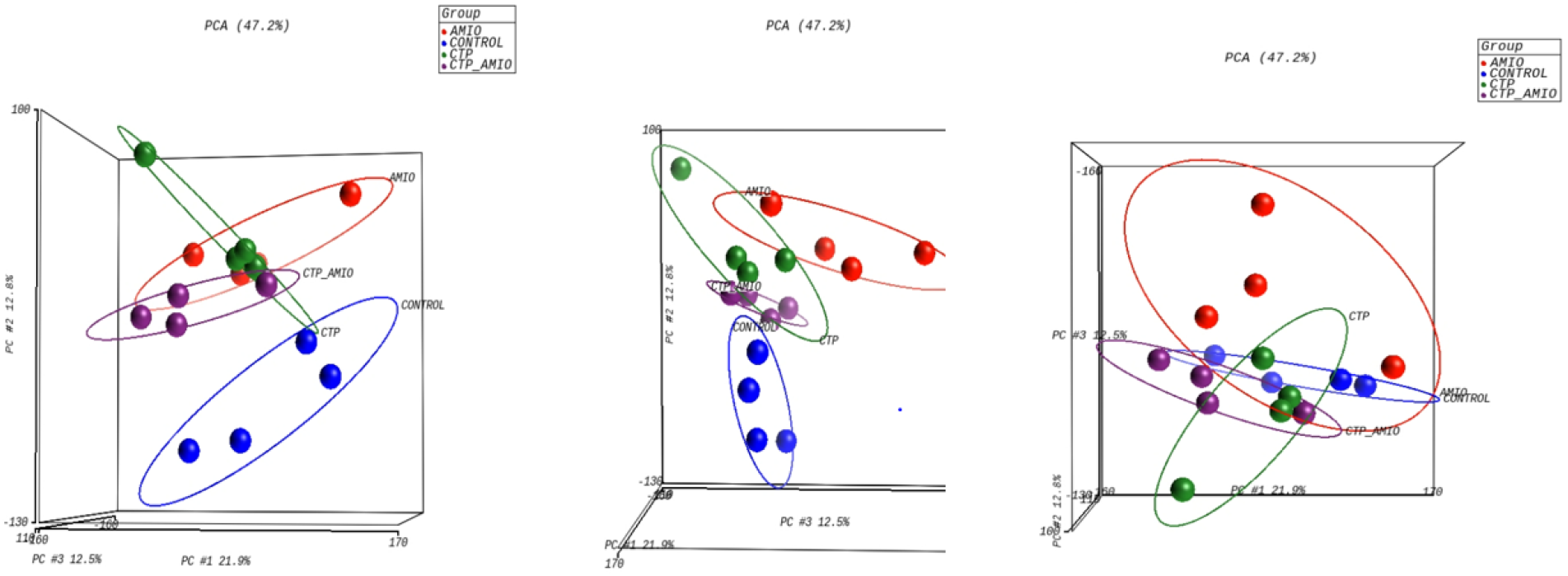
Principal component analysis showing reasonable clustering within groups and separation between different treatment groups.

**Figure 6:**
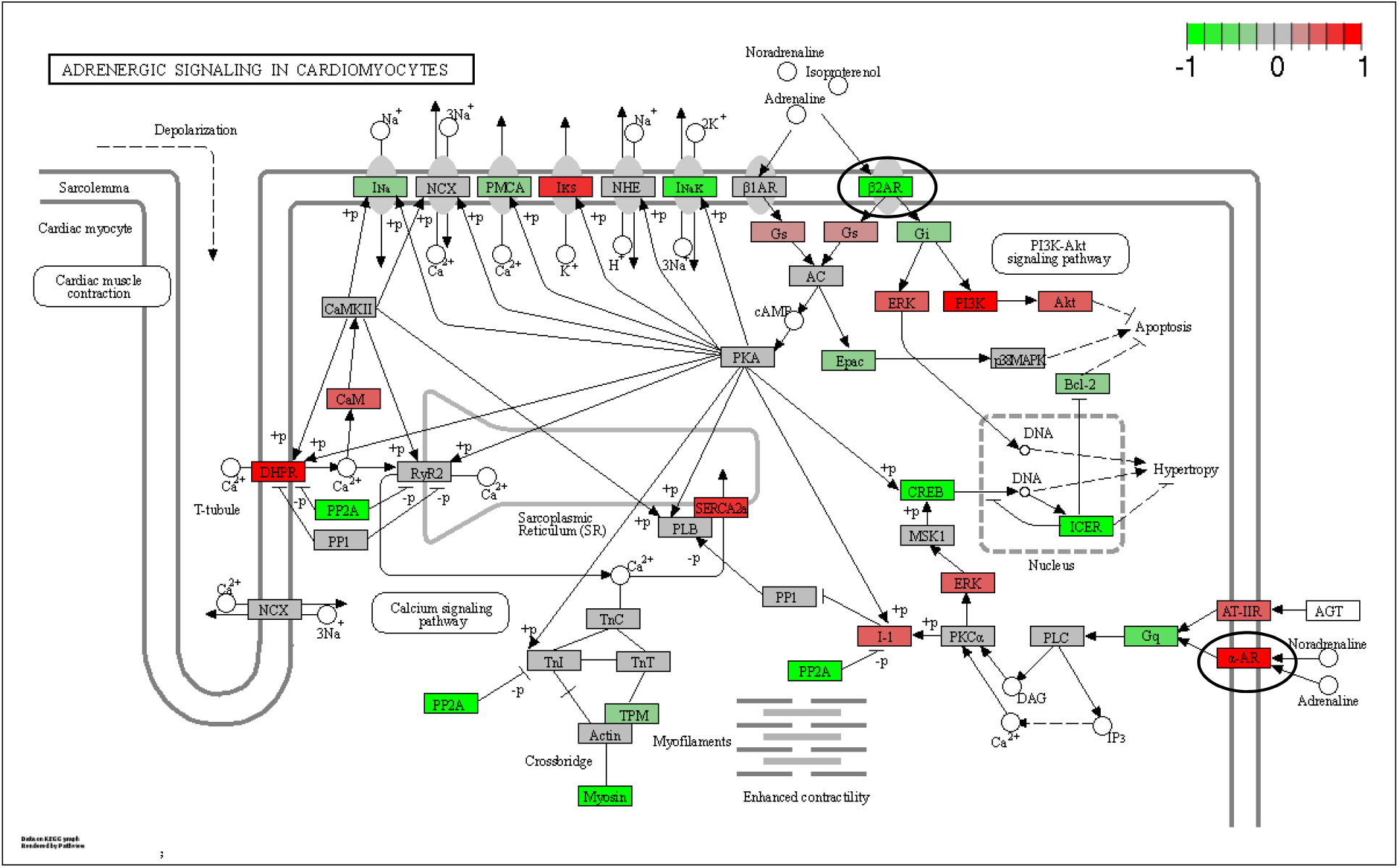
Differentially expressed genes in CTP treated guinea pig hearts compared with control hearts. β2-adrenergic receptor is significantly down-regulated, whereas α-adrenergic, receptor and calcium handling genes (*DHPR* and *SERCA2a*) are significantly upregulated (all circled in black).

**Figure 7:**
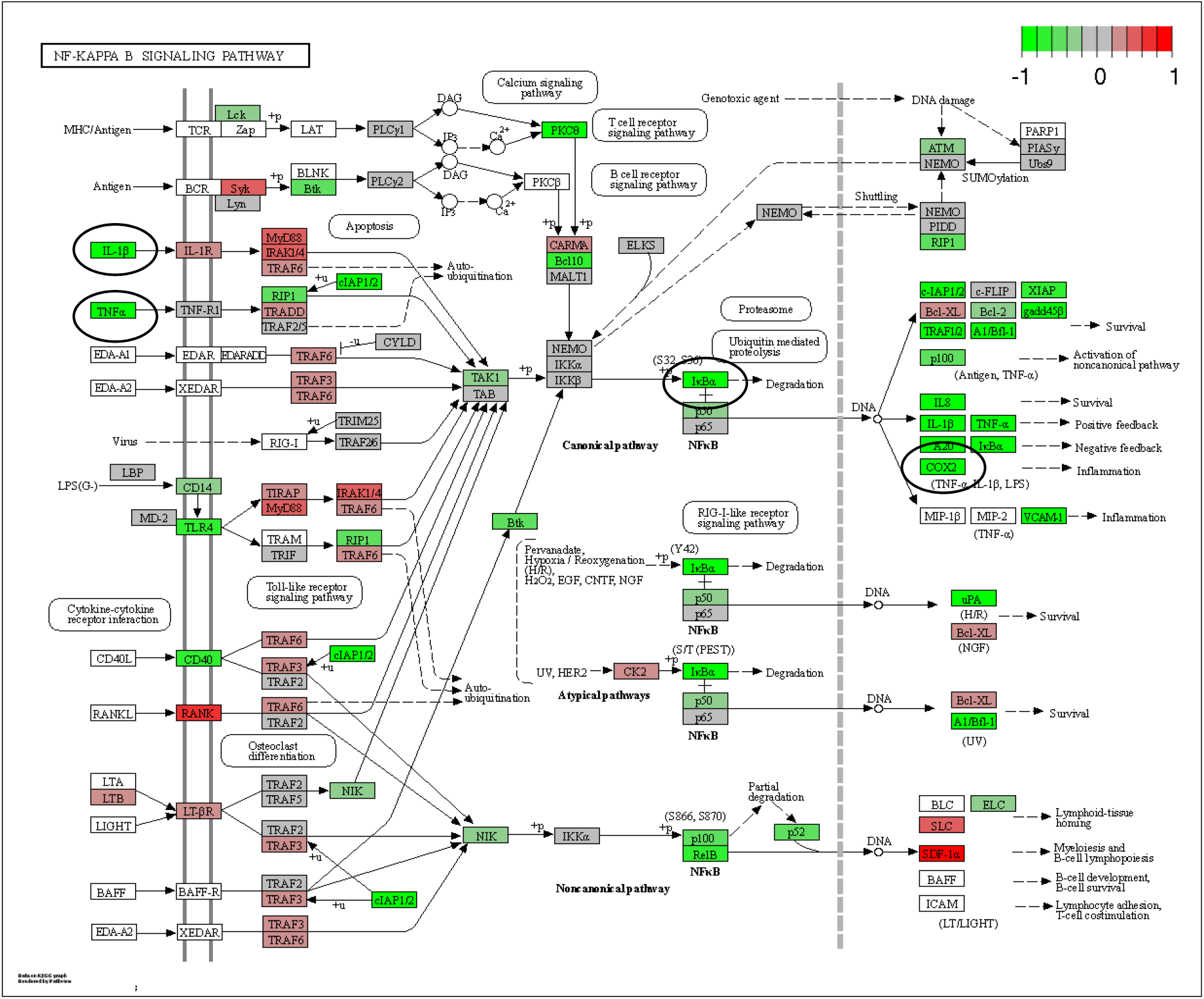
Differentially expressed genes (DEGs) in CTP treated guinea pig hearts compared with control hearts. Pro-inflammatory genes in the NF-κB pathway (*IκBα*), as well as *TNF-α, IL-1β*, and *COX2* are significantly down-regulated (all circled in black).

## Discussion

The targeted delivery of therapeutics to the heart remains an elusive goal but one of high clinical impact. In the current body of work, we show the chemical possibility of conjugating amiodarone to a novel, synthetic, non-naturally occurring, cardiomyocyte-targeting peptide or CTP. The conjugation was through a covalent linker containing a disulfide bond which was stable at 37°C for upto 22 days. Uptake of this conjugate by GP hearts was indirectly evidenced by decrease in voltage and calcium conduciton velocities similar to amiodarone alone, but at ∼1/15^th^ the molar equivalent total dose. CTP-amio had additional effects on the GP hearts, like an increase in resting heart rates, decrease in APand CaT durations which were not caused by amiodarone treatment alone. These effects appear to be driven by the CTP portion of the conjugate and were seen with CTP alone injections, to a similar extent. These effects of CTP could potentially be due to changes in adrenergic receptor and calcium handling protein expression, as seen in our RNA sequencing experiments. A surprise finding was an anti-inflammatory effect of CTP alone compared to control hearts. There are multiple literature reports, in cells other than cardiomyocytes, showing that a decrease in cytosolic calcium leads to inhibition of NF-κB activation, which is a transcription factor upstream of TNF-α and IL-1β^38-42^. Based on our data and the existing literature, we hypothesize that CTP increases expression of calcium handling genes leading to decrease in free cytosolic calcium and decrease in NF-κB expression leading to an anti-inflammatory effect.

Cell penetrating peptides have been studied intensively as vectors for the last 25 years. There is emerging evidence that they by themselves may possess anti-bacterial or anti-tumor efficacy. However, this observation has been attributed largely to disruption of cell membranes expressing high levels of certain phospolipids or sphingolipids characteristic of tumor cells. We had considered CTP to be an inert, cardiomyocyte-targeting peptide. CTP having salutary effects on calcium handling and supressing expression of pro-inflammatory genes was an unexpected finding. Whether this is a dose-dependent effect seen as a result of fairly high and repetitive dosing of CTP remains to be seen.

CTP has been shown to be cardiomyocyte specific by independent investigators. In their work, Avula and colleagues showed that placing CTP on a pegylated nanoparticle bearing photosensitizer led to cardiomyocyte-specific ablation with sparing of “innocent bystander” cells like myofibroblasts and endothelial cells, similar to our findings^28^. Additionally, exosomes labeled with CTP and loaded with anti-RAGE (*R*eceptor for *A*dvanced *G*lycation *E*nd-products) siRNA was able to ameliorate myocarditis in a rat model^30^. Although our initial phage display work that led to the identificaiton of CTP was carried out in a rat cardiomyoblast cell line, with subsequent cycles of *in vivo* phage display in mice^26^, CTP is not species limited to mice, as its efficacy as a vector has been shown in sheep^28^ and rat models^30^ for delivery of a myriad of cargoes. Indeed, our own lab showed that human heart tissue explanted from patients undergoing heart transplants could be successfully transduced with fluorescently labeled CTP with uptake of the peptide by normal cardiomyocytes while sparing the fibroblasts making up the scar tissue. Cardiomyocyte uptake of CTP was not simply due to an increase in plasma membrane permeability, as demonstrated by a lack of Evans blue uptake^31^

Multiple investigators have tried to target the delivery of amiodarone to the heart. One approach has been to apply amiodarone directly to the epicardium using amiodarone-infused hydrogels^43-45^ or via atrial patches^46^. This may be of utility under certain circumstances like patients undergoing coronary artery bypass grafting or valve replacements, but are not practical for routine, long-term treatment. Amiodarone loaded cyclodextrin nanoparticles efficiently increase cardiac uptake of amiodarone in macrophage abundant tissues such as in the inflamed heart in the setting of myocarditis, resulting in decreased severity of off-target systemic toxicity, but it does not solely target the cardiomyocytes of the non-inflamed heart^47^. In a similar manner, amiodarone loaded Poly(lactic-co-glycolic acid nanoparticles^48^ or liposomal based amiodarone formulations^49,50^ increase solubility of the lipophilic amiodarone drug resulting in controlled release which reduces systemic toxicity but are not exclusive to the targeting of cardiac tissues. Our approach would have targeted delivery specifically to cardiomyocytes for arrhythmia management.

Our work has raised several interesting questions. The study utilized fairly high doses of amiodarone in GPs (80mg/Kg) equivalent almost to 10-fold the human loading dose of 400mg twice daily. Whether CTP will still alter gene expression at lower doses or “maintenance” doses remains to be seen. Our work generated several hypotheses linking CTP to changes in calcium handling and subsequent decrease in NF-κB activation, which are currently under active study. Additionally, the efficacy of CTP-amio in a relevant animal model of afib remains to be seen.

## Acknowledgments

The authors would like to thank the University of Pittsburgh RNA sequencing facility (William McDonald, Rania Elbakri) for sequencing the mRNA from guinea pig hearts, and Mayo Clinic bioinformatics core (Asha Nair) for analyzing the RNA sequencing data.

